# UL135 and UL136 Epistasis Controls Reactivation of Human Cytomegalovirus

**DOI:** 10.1101/2023.01.24.525282

**Authors:** Melissa A. Moy, Donna Collins-McMillen, Lindsey Crawford, Christopher Parkins, Sebastian Zeltzer, Katie Caviness, Patrizia Caposio, Felicia Goodrum

## Abstract

Human cytomegalovirus (HCMV) is beta herpesvirus that persists indefinitely in the human host through a protracted, latent infection. The polycistronic *UL133-UL138* gene locus of HCMV encodes genes regulating latency and reactivation. While UL138 is pro-latency, restricting virus replication in CD34+ hematopoietic progenitor cells (HPCs), UL135 overcomes this restriction for reactivation. By contrast, UL136 is expressed with later kinetics and encodes multiple protein isoforms with differential roles in latency and reactivation. Like UL135, the largest UL136 isoform, UL136p33, is required for reactivation from latency in hematopoietic cells. Furthermore, UL136p33 is unstable, and its instability is important for the establishment of latency and sufficient accumulation of UL136p33 is a checkpoint for reactivation. We hypothesized that stabilizing UL136p33 might overcome the requirement of UL135 for reactivation. To test this, we generated recombinant viruses lacking UL135 that expressed a stabilized variant of UL136p33. Stabilizing UL136p33 did not impact replication of the UL135-mutant virus in fibroblasts. However, in the context of infection in hematopoietic cells, stabilization of UL136p33 strikingly compensated for the loss of *UL135,* resulting in increased replication in CD34+ HPCs and in humanized NOD-*scid* IL2Rγ_c_^null^ (NSG) mice. This finding suggests that while UL135 is essential for reactivation, it functions at steps preceding the accumulation of UL136p33 and that stabilized expression of UL136p33 largely overcomes the requirement for UL135 in reactivation. Taken together, our genetic evidence indicates an epistatic relationship between UL136p33 and UL135 whereby UL135 may initiate events early in reactivation that will result in the accumulation of UL136p33 to a threshold required for productive reactivation.

**SIGNIFICANCE:** Human cytomegalovirus (HCMV) is one of nine human herpesviruses and a significant human pathogen. While HCMV establishes a life-long latent infection that is typically asymptomatic in healthy individuals, its reactivation from latency can have devastating consequences in the immune compromised. Defining virus-host and virus-virus interactions important for HCMV latency, reactivation and replication is critical to defining the molecular basis of latent and replicative states and in controlling infection and CMV disease. Here we define a genetic relationship between two viral genes in controlling virus reactivation from latency using primary human hematopoietic progenitor cell and humanized mouse models.

## INTRODUCTION

The mechanisms by which a virus persists in a latent state for a protracted period and the events that trigger reactivation are poorly understood. DNA herpesviruses are a robust model for elucidating mechanisms of viral latency because of the complex virus-host interactions that allow the virus to “sense” and “respond” to changes in host biology. Human cytomegalovirus (HCMV) is a large ∼236-kb double stranded DNA beta herpesvirus with a global seroprevalence of approximately 60% to 90% based on geographic and socioeconomic factors (1–3). Infection is typically asymptomatic in immunocompetent individuals. Once latency is established, subclinical sporadic reactivation and virus-shedding can occur throughout the lifetime of the host (4, 5). For immunocompromised individuals with inadequate cellular immunity, reactivation from latency can result in morbidity and mortality, particularly in stem cell and solid-organ transplant recipients (4–8). Further, congenital CMV infection can cause significant neurodevelopmental abnormalities, affecting approximately 1 in 150 babies born in the United States (9–15). HCMV infection may enhance immune responses in young adults but may be a driver in age-related pathologies and altered T cell homeostasis (16–18). Therefore, understanding the viral and host mechanisms underpinning HCMV persistence and reactivation is critical to developing strategies to control its reactivation and associated pathologies.

HCMV exhibits broad intra-host tropism. HCMV replicates productively in human primary fibroblasts, but also in epithelial and endothelial cells. HCMV latency has been best characterized in hematopoietic progenitor cells (CD34+ HPCs) and cells of the myeloid lineage (19). During latency, viral genomes are maintained, viral gene expression is restricted, and no new viral progeny are made. HCMV reactivates in response to viral and host cues including cellular stress, inhibition of PI3K/Akt signaling, differentiation, or steroid treatment to re-initiate viral gene expression (4, 20–26). Viruses have evolved complex gene networks to sense and respond to multiple environmental stimuli that can feedback to impact viral gene expression (19, 27, 28).

The ∼15-kb UL*b’* region of the HCMV genome spans *UL133* to *UL154* and encodes genes important to the regulation of immune responses, viral persistence, and dissemination (4, 29–33). The UL*b’* region is present in low-passage strains and clinical isolates but lost during serial passage of the virus in fibroblasts, thus partly or entirely lacking in most laboratory-adapted strains. While dispensable for replication in fibroblasts, these genes undoubtedly play important roles in other contexts of infection in the host (34–37). We have characterized a 3.6-kb polycistronic gene locus within the UL*b*’ region encoding four genes, *UL133, UL135, UL136,* and *UL138,* collectively referred to as the *UL133-UL138* locus (19, 38). On whole, the *UL133-UL138* locus is suppressive to viral replication (34, 35). Using recombinant viruses containing disruptions of a single gene or combinations of genes in this locus, we have defined genetic phenotypes for each gene with respect to the establishment of or reactivation from latency in hematopoietic cells. UL138 is pro-latency in that genetic disruption of *UL138* translation allows HCMV to replicate in the absence of a reactivation stimulus and results in increased viral yields relative to the parental WT virus (34, 39). Conversely, *UL135* is pro-replication and necessary for reactivation from latency (40). UL135 functions, in part, to overcome the suppressive effect of UL138 for reactivation. Both *UL135* and *UL138* are expressed early prior to the onset of viral DNA synthesis. *UL135* also has an epistatic relationship with the viral serine-threonine protein kinase, UL97, whereby UL135 function confers a heightened requirement for UL97 for viral DNA synthesis and viral gene expression in a productive infection (41). Although UL97 is the only HCMV-encoded kinase, it is remarkably dispensable for replication in the laboratory-adapted strains lacking UL135 (41, 42).

*UL136* is a complex gene that encodes five protein isoforms resulting from alternative transcription initiation start sites (43). UL136 protein isoforms have a hand in both latency and reactivation with differential early-late expression kinetics (43, 44). The UL136-33kDa (UL136p33) and UL136-26kDa (UL136p26) membrane-associated isoforms are required for reactivation, with phenotypes that mirror a loss of *UL135* when disrupted (44). The UL136-25kDa (UL136p25) isoform has context-dependent roles where disruption of this isoform results in a more replicative virus in HPCs infected in vitro, but a virus that fails to reactivate humanized mice (44). The UL136-23kDa (UL136p23) and UL136-19kDa (UL136p19) isoforms are soluble and required for latency, similarly to *UL138* in that when they are disrupted the virus replicates in hematopoietic cells in the absence of a reactivation stimulus (43, 44). In contrast to *UL135* and *UL138*, UL136 gene products depend on viral DNA synthesis for maximal accumulation (43). Given these expression kinetics and the differential roles in latency and reactivation of the UL136 isoforms, we postulate that UL136 functions to toggle the balance between a UL138-dominant latent state and a UL135-dominant reactivated state in response to host cues.

We have recently defined an important regulatory checkpoint for controlling accumulation of the UL136p33 isoform for reactivation. UL136p33 is unstable relative to other UL136 isoforms (45). We have further shown that the host E3 ubiquitin ligase, Inducible Degrader of Low-Density Lipoprotein Receptor (IDOL, also known as MYLIP) targets UL136p33 for rapid proteasomal degradation. When UL136p33 is stabilized, HCMV is more replicative in the absence of a reactivation stimulus in CD34+ HPCs. IDOL is highly expressed in hematopoietic cells, but down regulated sharply upon differentiation. Further, induction of IDOL during infection in CD34+ HPCs restricts replication, whereas knockdown of IDOL increases virus gene expression in hematopoietic models. This work demonstrates the importance of UL136p33 instability for the establishment of latency and defines a key cellular pathway regulating UL136p33 levels and HCMV fate decisions with regards to entry into and exit from latency (45).

The independent requirements for UL135 and UL136p33 for reactivation from latency along with their sequential temporal expression pattern suggests an epistatic relationship between *UL135* and *UL136* genes for controlling latency and reactivation. We hypothesized that stabilizing UL136p33 might obviate the need for UL135 for reactivation. To explore the possible interdependence between UL135 and UL136p33 for reactivation, we generated UL136 recombinant viruses where UL136p33 was stabilized in the presence or absence of UL135. We demonstrate here that stabilizing UL136p33 does not compensate for the loss of *UL135* in fibroblasts. However, in a latent infection, stabilization of UL136p33 strikingly compensates for the loss of *UL135*, resulting in enhanced virus replication (a failure to establish latency) and genome amplification. Importantly, this phenotype is recapitulated in humanized NSG mice. Taken together, our evidence suggests an epistatic relationship between UL136p33 and UL135 for controlling HCMV reactivation from latency.

## RESULTS

### Stabilizing UL136p33 modestly alters viral gene expression in a productive infection

We have previously shown that UL135 exhibits early and UL136 isoforms exhibit early-to-late expression kinetics in a productive infection (43), that UL136p33 is unstable and its accumulation is required for reactivation (45). Further, as UL135 and UL136p33 are both required for latency, we hypothesized an epistatic relationship whereby UL135 function may drive aspects of infection required for the accumulation to UL136p33. If true, then stabilizing UL136p33 might overcome the requirement for UL135. UL136p33 is stabilized by the substitution of four lysine residues for arginine, *UL136_myc_*Δ*K*→*R.* To investigate the possible relationship between UL135 and UL136p33, we generated recombinant bacterial artificial chromosomes (BAC) by substituting *UL136* variants encoding the C-terminal myc epitope-tagged version of UL136, *UL136*_myc_, or the stabilized version of UL136, *UL136_myc_*Δ*K*→*R*, into a previously characterized *ΔUL135_STOP_* BAC where *UL135* expression is disrupted by the insertion of stop codons (43, 45) . The myc epitope tag is required to detect UL136 as we have been unsuccessful in generating antibodies to UL136. Further, while UL136 is important for infection in hematopoietic cells, UL136 is expressed at such low levels in CD34+ HPCs that it is undetectable even with the myc epitope tag (43, 44). A schematic of the recombinant viruses used in this study is shown in (Fig. 1A). Whole genome sequencing was performed to ensure mutations were located at the desired positions on the viral genome and that recombination did not affect other regions of the genome (Fig. 1B). In addition, a schematic of the specific nucleotides that were mutated are specified for each recombinant virus (Fig. 1C).

**Fig. 1.**
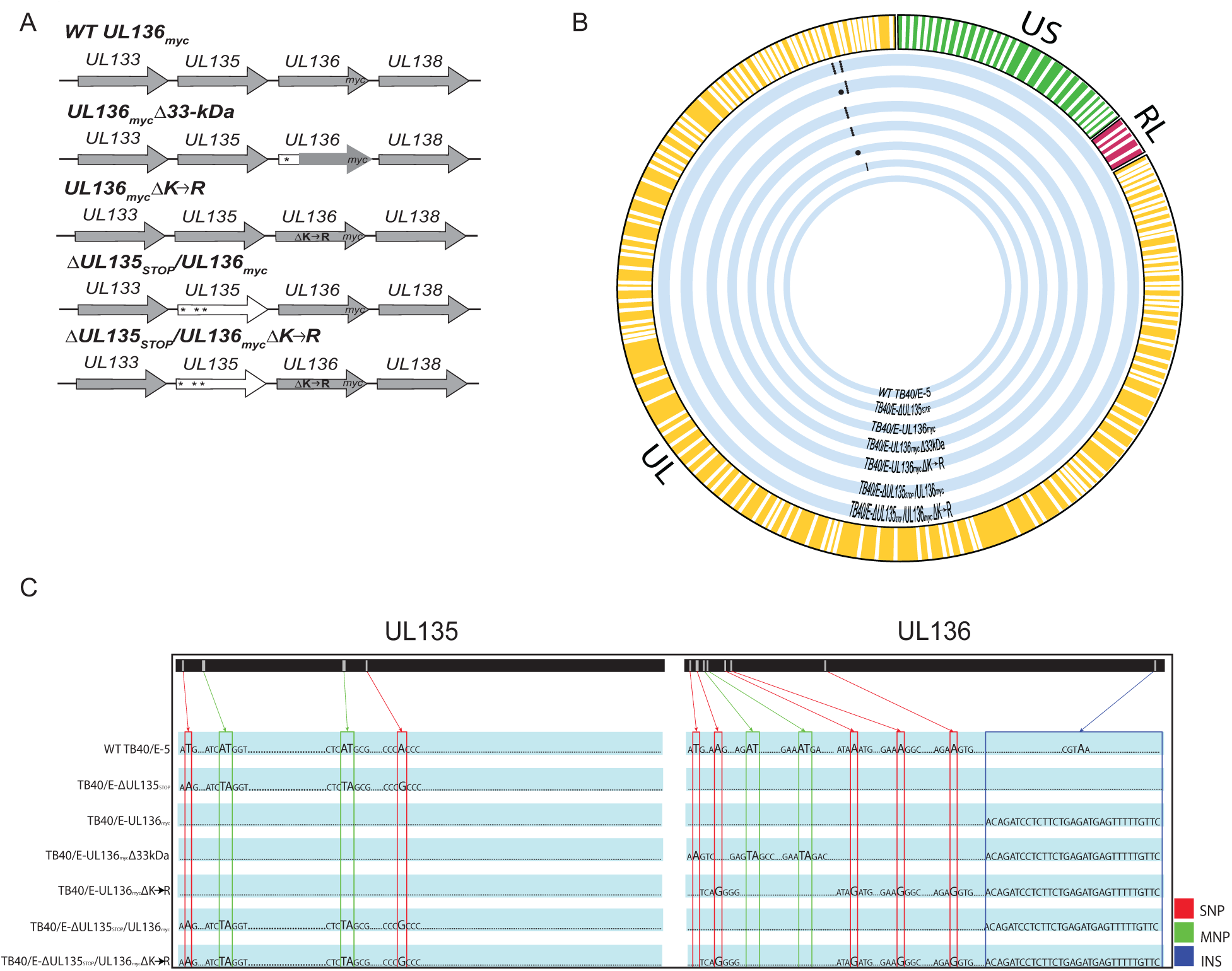
Schematic and whole genome sequencing of viruses used in this study. (A) Schematic of changes within the UL133-UL138 locus in TB40/E recombinant viruses indicated. Parental (WT) *UL136_myc_* expresses UL136 fused in frame with a C-terminal *myc* epitope tag. All recombinant viruses used in this study are derived from this virus and express the myc-epitope tagged version of *UL136* for protein detection. *UL136_myc_*D*33kDa* contains stop codon substitutions for 5’ AUGs to disrupt expression of UL136p33. *UL136_my_*Δ*K*→*R* contains arginine substitutions for all four lysine residues in UL136 (amino acid positions 4, 20, 25, and 113). D*UL135_STOP_/*D*UL136_myc_* contains stop codon substitutions for 5’ AUGs to abrogate synthesis of UL135 protein (amino acid positions 1, 27, and 97). D*UL135_STOP_/*D*UL136_myc_*Δ*K*→*R* combines the stop codon substitutions in UL135 with the lysine to arginine substitutions in UL136 described above. *, Indicates stop codon substitutions for methionine codons. (B) Whole genome sequencing and alignment of parental (TB40/E) and recombinant HCMV Bacterial Artificial Chromosomes (BACs) was performed to identify alterations in each recombinant virus. Thick blue lines represent each recombinant virus as labeled and mutations in each genome are denoted by black dots. (C) Zoomed in pictograph of *UL135* and *UL136* genes with specific nucleotide changes evident in sequencing. Single nucleotide polymorphisms (SNPs) are boxed in red; multiple nucleotide polymorphisms (MNPs) are boxed in green; insertions (INS) are boxed in blue. Mutations for each recombinant virus are compared back to the WT TB40/E-5 parental virus.

We first characterized viral gene expression during infection in MRC5 fibroblasts. Fibroblasts were infected with *UL136*_myc,_ *UL136_myc_*Δ*K*→*RR*, D*UL135_STOP_/UL136_myc_*, or D*UL135_STOP_/UL136_myc_*Δ*K*→*R* at an MOI of 1 and viral protein accumulation was analyzed over a time course (Fig. 2A). As expected, UL135 was not detected in D*UL135_STOP_/UL136_myc_* and D*UL135_STOP_/UL136_myc_*Δ*K*→*R* infections. Further, as previously reported, at a high MOI, representative immediate early, early, and late (IE1/2, UL44, pp28, and pp150 respectively) viral gene expression follows the standard gene expression cascade in the absence of *UL135* (40). Stabilizing UL136p33 (*UL136*_myc_Δ*K*→*R*) resulted in increased p33 protein levels, but decreased levels of middle UL136 isoforms, particularly p25, relative to *UL136*_myc_ infection (Fig. 2A-B). Further, stabilizing UL136p33 had little to no effect on early or late protein expression, represented by UL44 and pp150 respectively, relative to *UL136*_myc_ infection (Fig. 2A-B). We detect no significant differences in IE or early gene expression between the mutant viruses. Although, as previously reported (40), viruses lacking *UL135* may have modestly diminishes levels of IE or early proteins. Levels of the pp150 late protein were diminished in viruses lacking *UL135* and this was not rescued by the stabilization of UL136p33. With respect to the UL136 isoforms, UL136p33 levels were elevated in *UL136*_myc_Δ*K*→*R* infection, but only when *UL135* was present (Fig. 2A and C). We previously described the diminished accumulation of middle UL136 isoforms during infection with UL136_myc_Δ*K*→*R* (45). These results suggest that UL135 is necessary for maximal accumulation of UL136p33 at late times, while increased expression of UL136p33 diminishes middle UL136 isoform accumulation, some of which are pro-latency (43, 44).

**Fig. 2.**
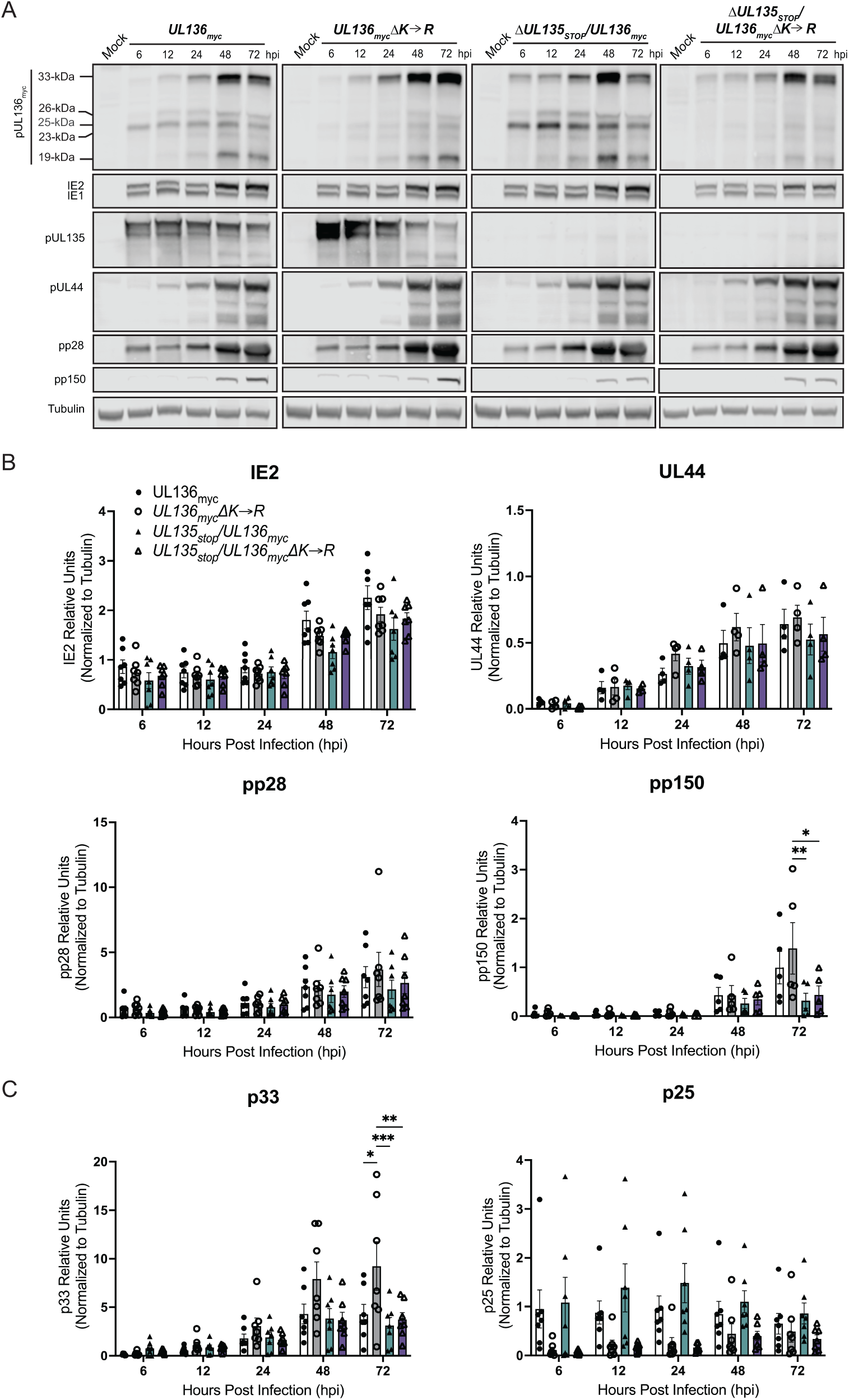
Stabilizing UL136p33 modestly alters viral gene expression in a productive infection. (A) MRC-5 lung fibroblast cells were infected with *UL136*_myc,_ *UL136_myc_*Δ*K*→*R*, D*UL135_STOP_/UL136_myc_*, or D*UL135_STOP_/UL136_myc_*D*K*à*R* recombinant viruses at MOI of 1. Lysates were collected over a time course of infection and immunoblotted using antibodies specific to the myc epitope tag (UL136 Isoforms), UL135 (UL135 isoforms), UL44, IE1&2 (clone 3H4), pp28 and pp150. Tubulin was used as a loading control. Representative blots are shown. (B-C) Protein levels quantified over multiple independent experiments using Image Studio Lite quantification software. Bars represent the averages of at least three independent experiments with standard deviation shown. Significance is calculated using two-way ANOVA with Tukey’s multiple comparisons test. *p<0.01; **p<0.001; ***p<0.0001.

### Stabilization of UL136p33 does not rescue viral yields associated with disruption of *UL135* in productive infection

We previously have shown that disruption of *UL136p33* alone or stabilization of UL136p33 does not affect virus yields resutling from a productive infection in fibroblasts (43, 45). Further, we have previously reported that disruption of UL135 results in a modest viral yield reduction in fibroblasts (40). To determine if stabilizing UL136p33 could rescue viral replication of Δ*UL135*_STOP_, we performed a multi-step growth curve (MOI 0.02) in fibroblasts infected with *UL136*_myc,_ *UL136_myc_*Δ*K*→*R*, D*UL135_STOP_/UL136_myc_*, or D*UL135_STOP_/UL136_my_*Δ*K*→*R* recombinant viruses (Fig. 3A). While *UL136_myc_*Δ*K*→*R* replicates with similar titers and yields as the parental virus, stabilization of UL136p33 did not compensate for the disruption of *UL135*; D*UL135_STOP_/UL136_myc_*Δ*K*→*R* replicated with similar kinetics and to similar yields as D*UL135_STOP_/UL136_myc_*.

**Fig. 3.**
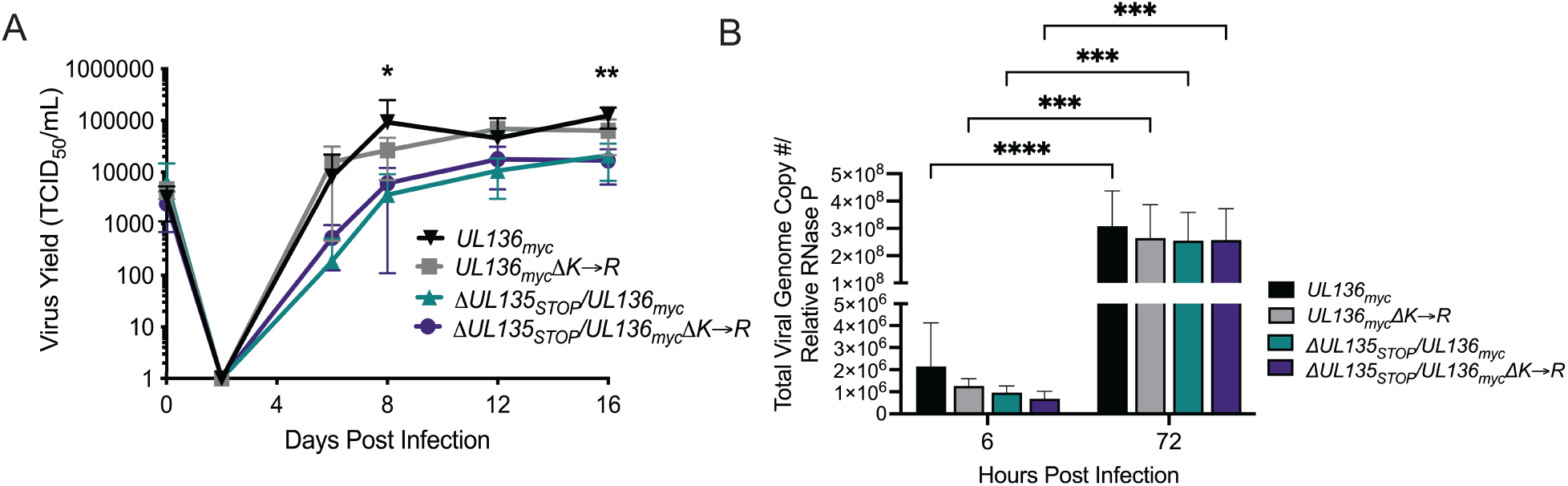
Stabilization of UL136p33 does not rescue viral yields associated with disruption of UL135. (A) Primary MRC-5 cells were infected with WT *UL136*_myc,_ *UL136_myc_*Δ*K*→*R*, D*UL135_STOP_/UL136_myc_*, or D*UL135 _STOP_/UL136_myc_*D*K*à*R* recombinant viruses and a multi-step (MOI of 0.02) growth curve was performed. Cells and culture supernatant were collected at the indicated timepoints, and virus titers were measured by TCID_50_. Data points represent the averages from three independent experiments and error bars represent standard deviations. Two-way ANOVA with Tukey’s multiple comparisons tests were performed to determine statistical significance for each mutant infection relative to the WT UL136_myc_ infection. *p<0.05 for WT *UL136_myc_* compared to *ΔUL135 _STOP_/UL136_myc_* and *ΔUL135 _STOP_/UL136_myc_*Δ*K*→*R* at 8 dpi and **p<0.01 for WT *UL136_myc_* compared to Δ*UL135 _STOP_/UL136_myc_* and *ΔUL135 _STOP_/UL136_myc_*Δ*K*→*R* at16 dpi. (B) Total DNA was isolated from primary MRC-5 cells infected at an MOI of 1 with WT *UL136*_myc,_ *UL136_myc_*Δ*K*→*R*, D*UL135 _STOP_/UL136_myc_*, or D*UL135 _STOP_/UL136_myc_*Δ*K*→*R* recombinant viruses at 6 and 72 hpi. The total number of viral genomes were quantified using qPCR with primers to β2.7kb relative to the cellular RNaseP gene. The viral genomes present in the cell at 6 hpi represent input genome copy number. Averages of duplicate measurements from three independent experiments with standard deviation bars are shown for each virus at each timepoint. Two-way ANOVA with Tukey’s multiple comparisons tests were performed to determine statistical significance values for each mutant infection relative to its 6 hpi time point (***, p<0.001 and ****, p≤0.0001).

We also analyzed viral genomes amplified in cells infected with WT *UL136*_myc,_ *UL136_myc_*Δ*K*→*R*, D*UL135_STOP_/UL136_myc_*, or D*UL135 _STOP_/UL136_myc_*Δ*K*→*R* recombinant viruses at an MOI of 1 (Fig. 3B). Total viral genomes were quantified using real time quantitative PCR (qPCR) with primers specific to sequences encoding the β2.7 region of the genome (46) at 6 and 72 hours post-infection (hpi). Viral genomes were amplified to similar levels in each infection, indicating that viral genome synthesis is not impacted by the loss of UL135 or the stabilization of UL136p33. The replication defect observed for Δ*UL135*_STOP_ infection in fibroblasts is due to late phase defects that were not rescued by the stabilization of UL136p33.

### Increased UL136p33 concentration does not enhance viral gene expression separate from viral DNA synthesis

Maximal expression of UL136 requires viral DNA synthesis (39, 40, 43). To examine the relationship between UL136 and viral gene expression around viral DNA synthesis, we analyzed the accumulation of viral proteins in fibroblasts infected with *UL136*_myc_ or *UL136_myc_*Δ*K*→*R* and treated with phosphonoacetic acid (PAA) to inhibit HCMV genome synthesis or a vehicle control (Fig. 4A). As previously shown, PAA treatment diminishes the accumulation of UL136 isoforms (43) and the canonical late protein, pp28, as well as amplification of IE2 at 48-72 hpi (43, 47, 48). While *UL136_myc_*Δ*K*→*R* increased levels of UL136p33, late-phase UL136p33 accumulation (48-72 hpi) is limited by PAA relative to the vehicle control (Fig. 4A). Quantification of UL136p33, UL135, IE2 and pp28 protein levels is shown in Fig. 4B. Stabilization of UL136p33 did not rescue late-phase IE2 or pp28 gene expression in infected cells where viral genome synthesis and entry into late phase was blocked by PAA (Fig. 4B). Further, stabilization of UL136p33 does not fully compensate for the onset of viral genome amplification and late phase events in driving maximal UL136p33 accumulation.

**Fig. 4.**
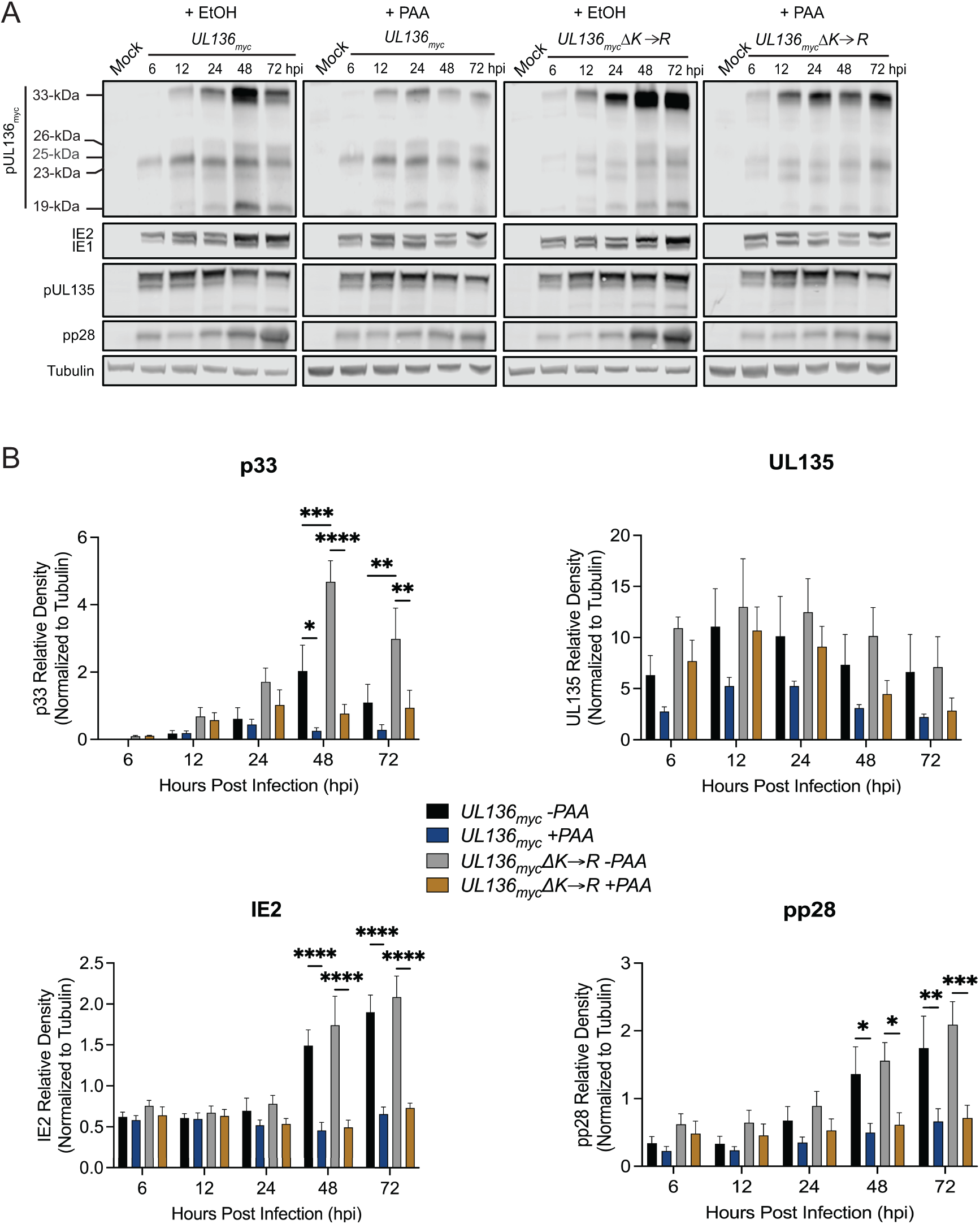
Increased UL136p33 concentration does not enhance viral gene expression separate from viral DNA synthesis. (A) MRC-5 cells were infected with WT *UL136*_myc_ or *UL136_myc_*D*K*à*R*, at an MOI of 1 and treated with PAA (add drug concentration) or ethanol as the vehicle control at the time of infection. Drug was re-freshed at 24 and 48 hpi to account for decay. Lysates were collected and immunoblotted at indicated time points for viral proteins using antibodies described in Fig. 1. Tubulin antibody was used as a loading control. Representative blots are shown. (B) Multiple independent experiments were quantified. Data points represent the averages from three independent experiments and error bars represent standard deviations. Two-way ANOVA with Tukey’s multiple comparisons tests were performed to determine statistical significance values for each infection relative to PAA treatment at each time point. *, p<0.05; **, p<0.01; ***, p<0.001; ****, p<0.0001.

### Stabilizing UL136p33 does not direct middle UL136 isoforms for proteasomal degradation

We have previously shown the UL136p33 is targeted for rapid proteasomal turnover by a host E3 ubiquitin ligase, IDOL (45). We next wanted to ask if the loss of middle UL136 isoforms when UL136p33 is stabilized (Fig. 2) is due to proteasomal degradation. To investigate this, we infected cells with either *UL136*_myc_ or *UL136_myc_*Δ*K*→*R* and treated cells with MG132 or vehicle 6 hours prior to collecting lysates at 24 hpi (Fig. 5A). All UL136 isoforms are quantified over multiple independent replicates in (Fig. 5B). As observed before, UL136p33 levels increased with MG132 treatment (45). Somewhat surprisingly, proteasomal inhibition also increased levels of UL136myc*K*→*R*, indicating that in addition to IDOL-dependent targeting to the proteasome, UL136 may also be targeted for ubiquitin-independent, proteasome-dependent turnover. Ubiquitin-independent, proteasome-dependent turnover has been reported for the pp71 HCMV tegument protein (49), as well as other viral or cellular oncoproteins (49–51). More work is required to fully investigate this possibility.

**Fig. 5.**
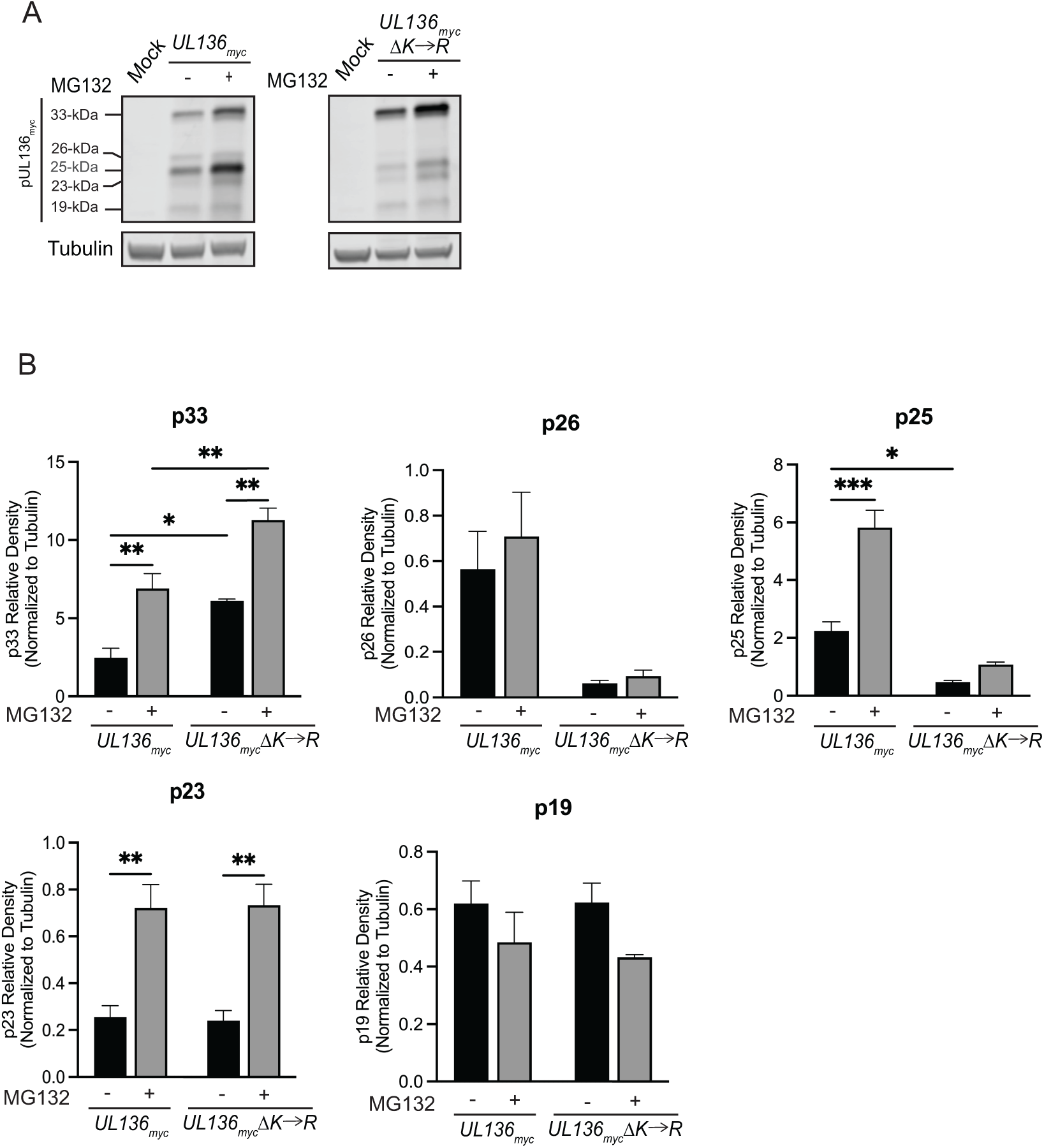
Stabilizing UL136p33 does not direct the smaller isoforms of UL136 for proteasomal degradation. MRC-5 fibroblasts were infected with either *UL136*_myc_ or *UL136_myc_*Δ*K*→*R*. 6 hours prior to lysate collection, infected cells were treated with 20 µM MG132 to inhibit the proteasome or vehicle control. Lysates were collected at 24 hpi and immunoblotted for myc to detect protein isoforms and tubulin (as a loading control) with antibodies described in Table 2. Three independent biological replicates were used to calculate statistical significance. Statistical significance was calculated through Two-way ANOVA with Tukey’s multiple comparisons for each isoform as follows. *, p<0.05; **, p<0.01; ***, p<0.001; ****, p<0.0001.

There are four lysine residues in UL136p33 (K4, K20, K25, and K113) and only K113 is shared by the UL136 isoforms smaller than p33. Like UL136p33, p25 and p23 are also rescued by MG132 treatment indicating that they are targeted to the proteosome. MG132 has no effect on p25 levels in the *K*→*R* mutant virus, suggesting that ubiquitination of K113 is required for its turnover. However, we have shown that IDOL does not target p25 (45) and so may be targeted by a distinct E3 ubiquitin ligase. UL136p23 expressed from the *K*→*R* mutant virus is rescued by MG132-treatment, indicating that it may be targeted for proteasome-dependent degradation, independently of ubiquitination at K113. UL136p26 levels were not affected by MG132, although UL136p26 levels were reduced in the *K*→*R* mutant virus infection, suggesting that this protein is not targeted for proteasomal degradation but that the stabilization of UL136p33 negatively affects the expression of p26. Lastly, UL136p19 was not affected by *K*→*R* substitution or by proteasomal inhibition, suggesting that UL136p19 is not regulated by the proteasome or UL136p33. Taken together, these results indicate that UL136p33 negatively impacts the expression of p26 and p25 through non-proteasomal degradation. Heightened UL136p33 levels likely downregulate transcription of smaller isoforms. However, differential transcriptional regulation of the UL136 isoforms is difficult to assess since UL136 isoforms are expressed from overlapping transcripts generated by alternative transcriptional start sites (43).

### Stabilizing UL136p33 rescues viral replication in the absence of *UL135* in CD34+ HPCs

Both *UL135* and *UL136p33* are required for reactivation from latency (40, 44). Further, stabilization of UL136p33 results in virus that is more replicative in CD34+ HPCs in the absence of a reactivation stimulus (45). We next wanted to determine if stabilizing UL136p33 could compensate for a loss of *UL135* in driving reactivation from latency in CD34+ HPCs. We infected CD34+ HPCs with WT *UL136_myc_*, *UL136_myc_*D*33kDa* (*TB40/E UL136_myc_* containing a stop codon disruption of the UL136p33 isoform) (44)*, UL136_myc_*Δ*K*→*R*, and D*UL135_STOP_/UL136_myc_*Δ*K*→*R* recombinant viruses. Infected HPCs (CD34+/GFP+) were purified by FACS and seeded into long-term bone marrow culture over a stromal cell support. At 10 dpi, the infected cell culture was split. Half the culture (live cells) was seeded by limiting dilution onto permissive fibroblast monolayers in a cytokine rich media to stimulate differentiation and reactivation (Reactivation). That other half of the culture was lysed and seeded in parallel on top permissive fibroblast monolayers by limiting dilution to determine infectious centers present prior to reactivation (Pre-Reactivation control) (52) (Fig. 6A). While we were unable to produce high enough titer stocks of the D*UL135_STOP_/UL136_myc_* virus sufficient for infection of CD34+ HPCs for all replicates, we have previously shown that Δ*UL135*_STOP_ viruses fail to reactivate, similarly to the *UL136_myc_*D*33kDa* infection (40, 53). As previously reported, the *UL136_myc_*Δ*K*→*R* infection resulted in a virus that replicated in CD34+ HPCs in the absence of a replication stimulus (45), indicating an inability to establish latency. Intriguingly, D*UL135_STOP_/UL136_myc_*Δ*K*→*R* infection resulted in a virus that was equally or more replicative than *UL136_myc_*Δ*K*→*R*. We also analyzed viral genome copy number at 10 dpi in two independent experiments (Fig. 6B). Consistent with the infectious centers measurements, viral genome levels were equivalent to or greater in the D*UL135_STOP_/UL136_myc_*Δ*K*→*R* infection relative to *UL136_myc_*Δ*K*→*R* infection. Taken together, these findings suggest that stabilization of UL136p33 can rescue the defect in reactivation associated with Δ*UL135*_STOP_ infection in CD34+ HPCs. This stands in contrast to the failure of *UL136_myc_*Δ*K*→*R* to rescue the defect in replication associated with disruption of *UL135* in fibroblasts (Fig. 3A) and suggests cell-type dependent functions or mechanisms for the *UL133/8* locus genes.

**Fig. 6.**
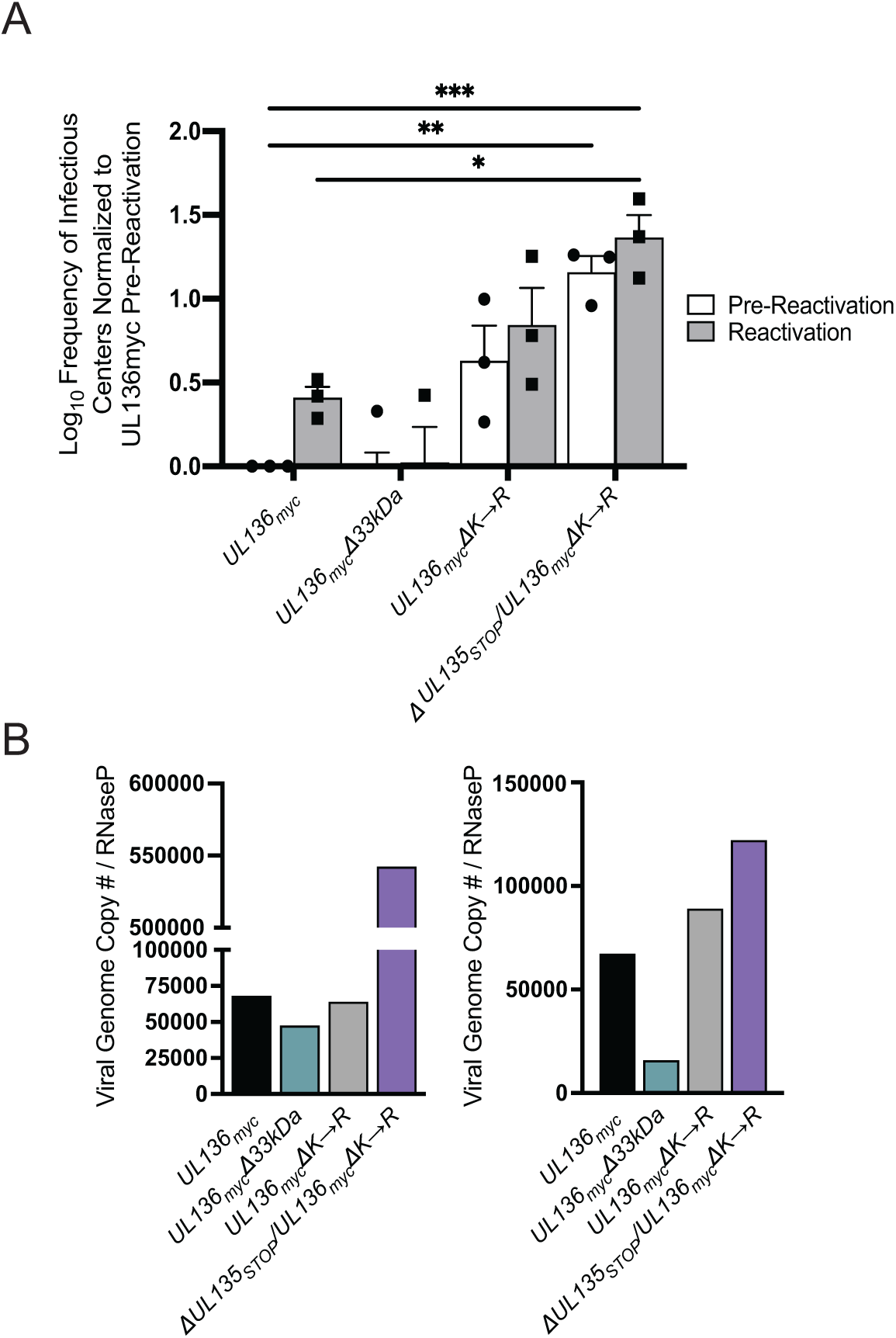
Stabilizing UL136p33 rescues viral replication in the absence of UL135 in CD34+ HPCs. (A) CD34+ HPCs were infected with WT *UL136_myc_*, *UL136_myc_*D*33kDa, UL136_myc_*Δ*K*→*R*, and D*UL135_STOP_/UL136_myc_*Δ*K*→*R* at an MOI of 2. At 24 hpi, CD34+/GFP+ (infected cells) were sorted and seeded into long-term bone marrow culture. After 10 days in culture, parallel populations of either mechanically lysed cells or whole cells were plated onto fibroblast monolayers in cytokine-rich media. 14 days later, GFP+ wells were scored, and frequency of infectious centers were determined by extreme limiting dilution analysis. The mechanically lysed population defines the quantity of virus present prior to reactivation (pre-reactivation; black bar). The whole-cell population defines the quantity of virus present after reactivation (reactivation; grey bar). The frequency was normalized to WT *UL136_myc_* pre-reactivation and the average of three independent experiments is shown. Statistical significance was calculated using two-way ANOVA with Tukey’s multiple comparisons test. *, *P*<0.05; **, *P*<0.01; ***, *P*<0.001). (B) Total DNA was isolated from CD34+ HPCs infected with WT *UL136_myc_*, *UL136_myc_*D*33kDa, UL136_myc_*D*K*à*R*, and D*UL135_STOP_/UL136_myc_*Δ*K*→*R* at an MOI of 2 on 10 dpi. The number of viral genomes relative to the level of RNaseP expression was quantified by qPCR using b2.7kb RNA gene- and RNaseP-specific primers. Two biological replicates from two independent cell donors are shown.

### Stabilizing UL136p33 compensates for the loss of *UL135* for viral replication in huNSG Mice

We next wanted to analyze the activity of these recombinant viruses in NOD-*scid* IL2Rγ_c_^null^ (huNSG) mice. We have previously demonstrated that UL136p33 is necessary for reactivation of HCMV post G-CSF stimulation in huNSG mice engrafted with CD34+ HPCs (44). While the huNSG mouse model has typically recapitulated results from our in vitro CD34+ HPC experimental latency model, the one notable exception is for the *UL136*_myc_*Δ25kDa* virus, which fails to reactivate in huNSG mice, but is more replicative in CD34+ HPCs infected in vitro (44). huNSG mice were sublethally irradiated and engrafted with human CD34^+^ HPCs. After CD34^+^ engraftment, mice were injected with human fibroblasts infected with WT *UL136_myc_*, *UL136_myc_*D*33kDa, UL136_myc_*Δ*K*→*R*, and D*UL135_STOP_/UL136_myc_*Δ*K*→*R* recombinant viruses. At 4 weeks post-infection, 5 mice from each group of 10 were treated with G-CSF and AMD-3100 to induce stem cell mobilization and HCMV reactivation. At 1-week post-mobilization, viral genome load was assessed in spleen and liver tissues from treated and untreated mouse groups to evaluate amplification of viral genomes and dissemination of infected cells to organs, as described previously (44).

As previously demonstrated, G-CSF mobilization of *UL136*_myc_-infected humanized mice resulted in increased viral genomes detected in the spleen and liver relative to unmobilized mice, consistent with reactivation of virus replication (Fig. 7). By contrast, *UL136*_myc_Δ33kDa failed to reactivate and similar levels of genomes were measured in G-CSF/AMD-3100-treated and untreated huNSG mice infected with either *UL136*_myc_ or *UL136_myc_*D*33kDa*. The viral genome copy number in the unmobilized and mobilized mice infected with *UL136_myc_*Δ*K*→*R* was equivalent to the G-CSF/AMD-3100-mobilized *UL136*_myc_ infection, indicating a failure to establish latency (replication in the absence of a stimulus). This phenotype recapitulates the *UL136_myc_*Δ*K*→*R* phenotype in CD34+ HPCs infected *in vitro* (Fig. 6). Strikingly, the D*UL135_STOP_/UL136_myc_*Δ*K*→*R* virus also recapitulated our *in vitro* studies with increased viral genome levels in the spleen and liver regardless of G-CSF/AMD-3100 treatment compared to mice infected with *UL136_myc_*Δ*K*→*R*. Taken together, these results support an epistatic relationship between UL135 and UL136 whereby stabilization of UL136p33 compensates for the loss of *UL135* in reactivation. Further, these data indicate a restrictive or deleterious consequence of UL135 function in reactivation and replication and that these defects can be overcome by robust expression of UL136p33.

**Fig. 7.**
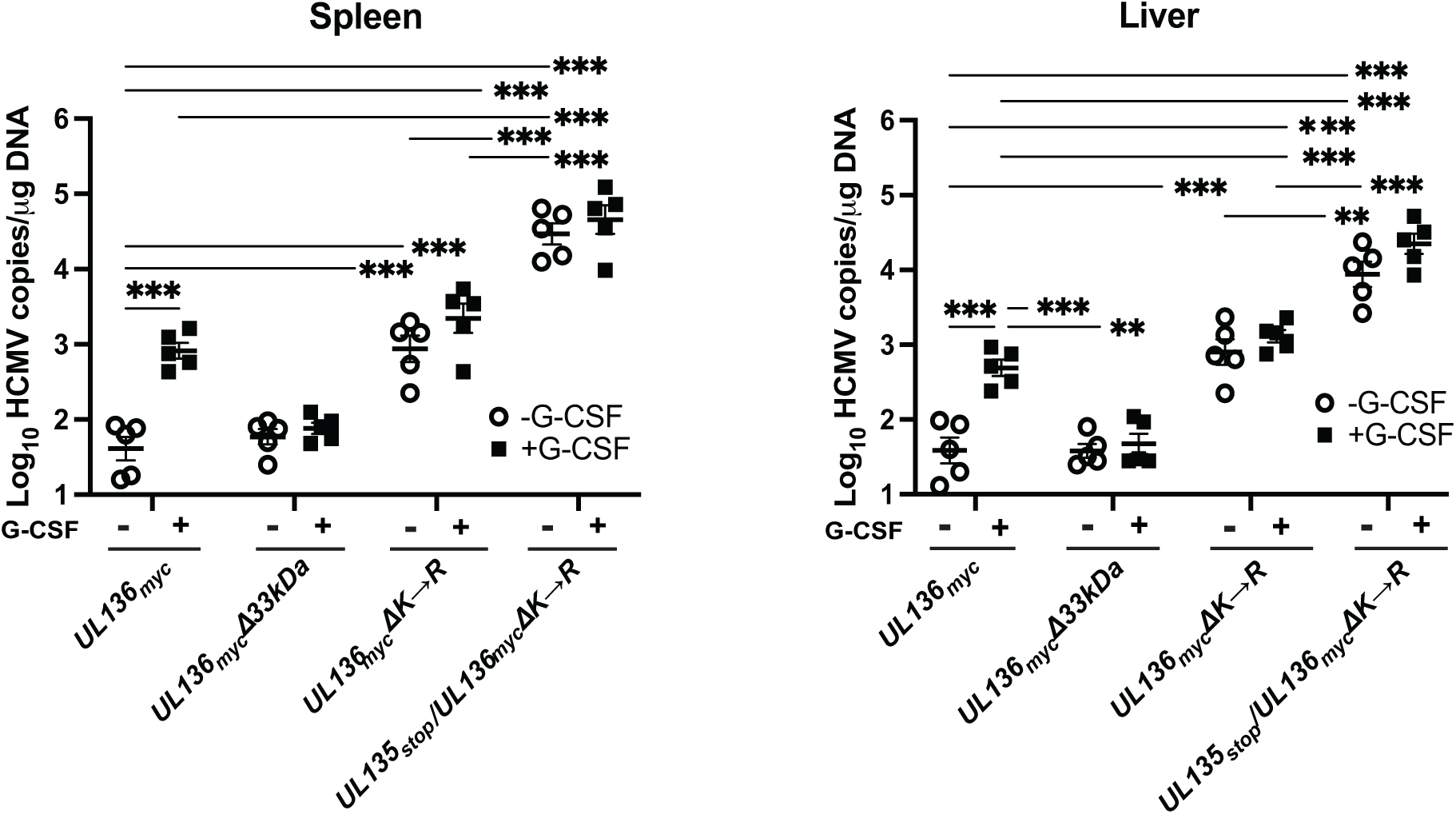
Stabilizing UL136p33 can compensate for a loss of UL135 for viral replication in huNSG Mice. Humanized NSG mice were injected with fibroblasts infected with either HCMV WT *UL136_myc_*, *UL136_myc_*D*33kDa, UL136_myc_*D*K*à*R*, and D*UL135_STOP_/UL136_myc_*Δ*K*→*R* (*n* = 10 per group). At 4 weeks post-infection, half of the mice were treated with G-CSF and AMD-3100 to induce cellular mobilization and promote HCMV reactivation. Control mice were left untreated. At 1-week post-mobilization, mice were euthanized, and tissues were collected. Total DNA was extracted using DNAzol, and HCMV viral load was determined by qPCR on 1 μg of total DNA prepared from spleen or liver tissue. Error bars represent standard error of the mean between average DNA copies from two or four tissue sections, respectively, for individual animals. All samples were compared by Two-way ANOVA with Tukey’s multiple comparisons test within experimental groups (nonmobilized [-G-CSF] versus mobilized [+G-CSF] for each virus and between all virus groups for both nonmobilized and mobilized conditions). *P*-values listed are for statistical significance where *P* < 0.01.

## DISCUSSION

Viruses use complex gene regulatory networks to coordinate major infection checkpoints. To direct the transition from latency to reactivation, herpesviruses must sense and respond to multiple cellular processes to exit the latent state for replication. HCMV has a large coding capacity at more than 170 genes for regulating infection in multiple cell types with the possibility of a much-expanded array of open reading frames (54). Many genes encode multiple proteins (referred to as isoforms), as is the case for UL136, UL44, UL99, and UL112-113, through alternative transcriptional and/or translational start sites within a single gene (34, 43, 55–59). Other herpesviruses, such as Kaposi’s Sarcoma-Associated Herpesvirus (KSHV) and herpes simplex type-1 (HSV-1) also have genes that encode for multiple protein isoforms with differential spatiotemporal expression – ORF50 and UL12, respectively (60, 61). While the full potential of protein isoforms encoded by single genes in herpesviral genomes remains to be determined, genes encoding multiple proteins allow for co-regulation of effectors to dictate infection outcomes. Protein isoforms encoded from a single gene may have synergistic or opposing roles in dictating patterns of infection in a context-dependent manner (4). Further, epistatic interactions of viral genes in the same locus (or gene circuit) add to the complexity of HCMV biology. This work investigates the role of *UL136*, and specifically the instability of the UL136p33 isoform, and its epistatic relationship with UL135. This question is significant to understanding the complex, context-dependent interactions between virus-virus and virus-host factors that influence the “decision” to enter and maintain latency or to reactivate. These epistatic interactions and the timing by which they occur allow HCMV to sense changes in cell physiology to maintain the latent state or reactivate to produce viral progeny.

Both UL135 and UL136p33 are important for reactivation from latency (40, 44). UL135-mutant viruses exhibit a modest defect for replication and production of progeny virus in fibroblasts (34, 40), whereas disruption or stabilization of UL136p33 had no impact on replication in fibroblasts (43, 45). While stabilizing UL136p33 did not compensate for the loss of *UL135* in late gene expression or virus yields during infection in fibroblasts (Figs. 2-3), stabilization of UL136p33 strikingly compensated for the loss of *UL135* in hematopoietic cells infected in vitro or in humanized mice, resulting in increased virus replication in HPCs in the absence of a stimulus for reactivation (Fig. 6-7). This indicates distinct cell type-dependent roles for UL135 and UL136p33 for replication in CD34+ HPCs and huNSG mice. These findings suggest that while UL135 is important to the initiation of or commitment to reactivation, accumulation UL136p33 represents a subsequent threshold for reactivation. Once UL136p33 accumulation is ensured, in this case experimentally by lysine to arginine substitution, UL135 is dispensable for replication in experimental HPC models of latency.

In collaboration with the Kamil lab, we previously defined an epistatic relationship between UL135 and UL97 in promoting late-phase viral gene expression, viral DNA synthesis, and viral replication in a productive infection (41). In these studies, UL97, the only HCMV-encoded kinase, was dispensable for replication in a laboratory strain, but required for replication only in a low-passage strain expressing *UL13*5. This result suggested that UL135 function conferred a requirement for UL97 or, in other words, UL97 compensated for deleterious effects of UL135. Similarly, *UL135*_STOP_/*UL136_myc_*Δ*K*→*R* replicates with or without a reactivation stimulus to enhanced levels compared to the *UL136_myc_*Δ*K*→*R* virus, particularly in the humanized mouse model (Fig. 6 and 7). Together with the UL97/UL135 relationship, this study suggests that UL135 functions, while important for reactivation, pose deleterious consequences for virus replication and that the virus encodes other genes to mitigate those effects.

While UL135 is expressed with early kinetics, UL136 and UL97 are expressed with early/late or leaky late kinetics and their maximal accumulation requires viral DNA synthesis and entry into late phase (43, 62). These kinetics, taken together with the results of this study, suggest a model whereby latency is driven by UL138 and the turnover of UL136p33. Reactivation from latency coincides with the expression of UL135, which functions, in part, to overcome the suppressive effects of UL138 (40, 63) and by driving UL136 expression, specifically the accumulation of UL136p33. At the same time, UL97 expression may compensate for deleterious effects of UL135 (41). UL136p33 accumulation is further fortified by the initiation of viral DNA synthesis (Fig. 4), indicating a second checkpoint (following UL135 expression) in the commitment to reactivation and replication (Fig. 8). This model suggests a stepwise progression towards reactivation where thresholds must be met to pass checkpoints that would otherwise maintain latency.

**Fig. 8.**
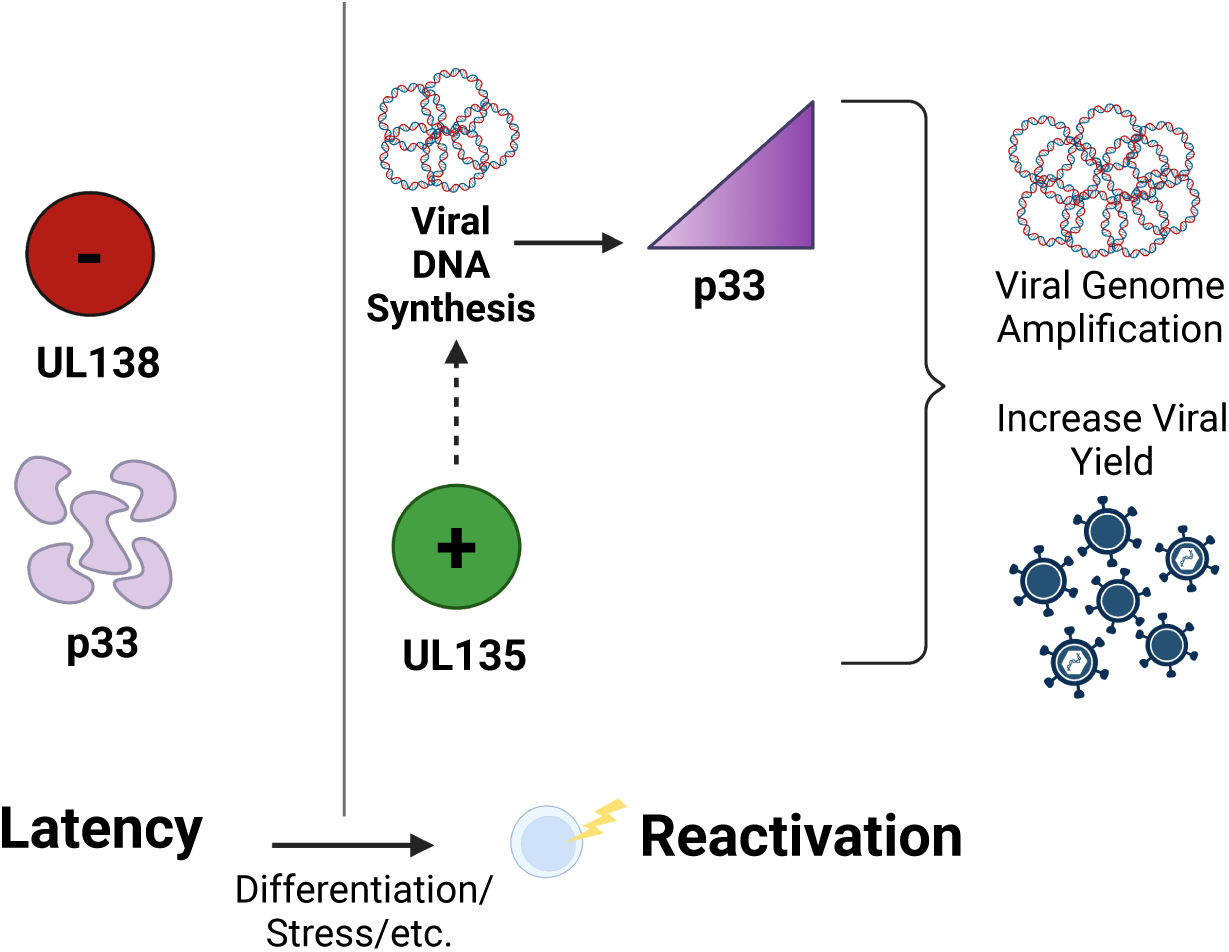
Model for UL135 and UL136p33 epistasis for coordinating reactivation of HCMV. We have previously shown that viral factors, such as UL138, function in a pro-latency manner in the establishment of latency. We have further demonstrated that the instability of UL136p33, driven by the IDOL E3 ligase, is also important for the establishment of latency. UL136p33 is required for reactivation. Reactivation cues stimulate increased expression of UL135 relative to UL138 and increased accumulation of UL136p33 through two mechanisms: (i) the loss of IDOL and (ii) increased UL136p33 synthesis following commitment to viral DNA synthesis. Because stabilization of UL136p33 precludes the need for UL135 for replication in HPCs, we propose the existence of checkpoints in the decision to reactivate where UL135 initiates reactivation and subsequent accumulation of UL136p33 is required to meet a threshold in the commitment to reactivation. Created with BioRender.com.

By this model, low levels of UL135 and UL136p33 are critical for the establishment of latency. We have previously shown that UL138 expression is driven by binding of the host EGR-1 transcription factor to sites just upstream of UL136 (64). As EGR-1 is highly expressed in HPCs, these cells are primed to express UL138 and the ratio of UL138 transcripts to UL135 transcripts that also encode UL136p33 is high. UL136p33 is maintained at low levels by the host E3 ubiquitin ligase, IDOL, which is also highly expressed in HPCs (45). Both EGR-1 and IDOL expression are sharply downregulated by differentiation (45, 64), a well-appreciated trigger of HCMV reactivation (4, 65). Therefore, differentiation is expected to increase the ratio of UL135 transcripts relative to UL138 transcripts (64) and increase expression and accumulation of UL136p33, which is further amplified by the onset of viral DNA synthesis and entry into late phase. Additional work is required to determine how UL135 promotes UL136p33 accumulation. Given the membrane association and cytoplasmic subcellular distribution of UL135 (40), it is unlikely that UL135 directly drives the transcription of UL136. However, *UL135* does encode a smaller isoform (initiation at methionine 97) that lacks the putative transmembrane domain and its role in infection has yet to be investigated (40). However, UL135 is well appreciated for its role in modulating host signaling and cytoskeletal organization and these roles could impact expression of UL136 indirectly (53, 63, 66).

Another potential area for viral control of reactivation is amongst the UL136 isoforms, as well as interaction between the UL136 isoforms and other UL133/8 locus genes. We have shown that while UL136p33 and p26 are important for reactivation, UL136p23/p19 are suppressive to replication for latency (44). Perhaps expression of UL136p33 overcomes the suppressive effects of the UL136p23/19 isoforms, similarly to the requirement of UL135 in overcoming UL138-mediated suppression (40). The antagonistic or synergistic roles of the UL136 protein isoforms may allow the virus to navigate the commitment to reactivation or the restoration of latency. The decision to maintain latency or reactivate at the point of viral DNA synthesis may depend on competition between or the differential accumulation of UL136 isoforms (44).

We have demonstrated here that stabilization of UL136p33 leads to a reduction in the protein levels of the middle UL136 isoforms (Fig. 2, 4 and 5), which may aid in allowing HCMV to commit to reactivation and replication. The differential regulation of UL136 isoform expression and their turnover by ubiquitin-dependent or -independent means is an important area for further investigation. UL136 isoforms are synthesized from unique transcripts derived from alternative transcriptional start sites and may be regulated in a context dependent manner (43). We have shown previously that HCMV reactivation from latency stimulates major immediate early gene expression from alternative promoter sequences embedded within intron A of the MIE transcriptional unit, while the MIEP apparently remains silent (47). In addition, Kaposi’s Sarcoma-Associated Herpesvirus (KSHV) *Orf50* encodes four different protein isoforms of RTA that temporally regulate each other’s expression for driving the viral gene expression cascade post-reactivation (60). This study demonstrated specifically that induction of the upstream *Orf50*, N5 promoter that produces RTA isoform 4, suppresses through transcriptional interference expression of RTA isoform 1. UL136p33 may be quelling the lower isoforms of UL136 through transcriptional interference of alternative promoters driving the smaller isoforms. Further work is required to define the promoters and transcription factors governing the UL136 isoforms. This will be imperative for understanding how their concentration and temporal expression kinetics regulate the transition between latency and reactivation of HCMV. Beyond transcriptional regulation, our findings presented here also indicate that some isoforms are targeted for proteasomal degradation (p33, p25, and p23), while others are not (p26 and p19). Of those targeted for proteasomal degradation, p23 appears to be targeted by ubiquitin-independent mechanisms since Δ*K*→*R* substitution had no effect on its turnover. Finally, the distinct mechanisms by which UL136 protein isoforms levels are regulated by degradation may reflect their distinct subcellular localization (44). The stabilization of p33 diminished levels of p26 and p25, neither of which could be rescued by proteasomal inhibition. This result likely reflects a negative feedback loop where elevated UL136p33 levels transcriptionally suppress the expression of middle isoforms from distinct transcripts (43).

This study highlights the complexity of the gene regulatory network controlling HCMV latency and reactivation. This gene regulatory network includes, but is not limited to, the UL133-UL138 locus. The ability of a virus to move between bistable infection states-latency or replication-requires that the virus can “sense” host cues, filter and utilize fluctuations (“noise”) to respond appropriately to the host cues to regulate infection fate decisions. This system is undoubtedly comprised of ultrasensitivity, positive and negative feedback mechanisms (67). UL136p33 may function in an ultrasensitive manner to control the switch for reactivation upon differentiation of HPCs (rapid accumulation of UL136p33 to a threshold to tip the system to reactivation). Going forward, it will be critical to understand the regulation of the HCMV gene regulatory network, particularly the possibility that UL136 isoforms are differentially regulated in response to host cues. Do UL136 isoforms help the virus stabilize or filter noise? Biological systems are comprised of intrinsic and extrinsic noise, and it is critical that a system can filter noise to respond appropriately; a virus that reactivates indiscriminately risks elimination. Intrinsic noise that may impact virus decisions include, nuclear architecture (68), nucleosome occupancy and positioning and epigenetic modification of chromatin (69, 70), TATA-box binding specificity (71–73), transcriptional pausing (74), and the regulation of protein levels through mRNA degradation, translation, and protein degradation (75, 76). Extrinsic noise includes availability of basic cellular machinery (77), pathway-specific propagation (78), micro-fluctuations in cellular environment (79, 80), cell division or asymmetric partitioning of cellular contents (81). HIV has been shown to attenuate noise through a non-transcriptional negative-feedback circuit where viral proteins feedback to deplete their own unspliced mRNA precursor to stabilize commitment decisions (82). It will be important to define these mechanisms regulating HCMV fate decisions and how UL136 and other viral genes contribute to fate decisions to understand viral control and pathogenesis.

## MATERIALS AND METHODS

### Cells

Human primary embryonic lung fibroblasts (MRC-5, purchased from ATCC; Manasas, Virginia) were cultured in Dulbecco’s modified Eagle’s medium (DMEM) and supplemented with 10% fetal bovine serum (FBS), 10mM HEPES, 1mM sodium pyruvate, 2mM L-alanyl-glutamine, 0.1mM non-essential amino acids 100 U/mL penicillin, and 100 μg/mL streptomycin. Institutional Review Board approved protocol to obtain human cells from bone marrow transplant waste at the University Medical Center at the University of Arizona. Specimens were de-identified and provided as completely anonymous samples. CD34^+^ HPCs were isolated and cultured as previously described (34, 52). Briefly, CD34^+^ HPCs were isolated using the CD34 MicroBead Kit (MACS, Miltenyi Biotec, San Diego, CA). Pure populations of CD34^+^ HPCs were subsequently cultured in MyeloCult H5100 (Stem Cell Technologies, Cambridge, MA) supplemented with hydrocortisone, 100 U/mL penicillin, and 100 µg/mL streptomycin and maintained in long-term co-culture with M2-10B4 and Sl/Sl murine stromal cells lines (kind gift from Stem Cell Technologies on behalf of D. Hogge, Terry Fox Laboratory, University of British Columbia, Vancouver, BC) (34, 52). All cells were maintained at 37°C with 5% CO_2_.

### Recombinant viruses

The TB40/E BAC was engineered to express the green fluorescent protein (GFP) driven by the SV40 early promoter located in the intergenic region in between US34 and TRS1 genes as a marker for infection (34, 83). All recombinant viruses were created using a two-step, positive/negative selection approach that leaves no trace of the recombination process (39, 84, 85). Three recombinant viruses used in this study were previously engineered – *UL136*_myc,_ *UL136_myc_*D*33kDa,* and *UL136_myc_*Δ*K*→*R* as described previously (43, 45). Two new recombinant viruses were generated for this study - D*UL135 _STOP_/UL136_myc_* (to detect the UL136 isoforms when *UL135* is disrupted) and D*UL135 _STOP_/UL136_myc_*Δ*K*→*R* (where UL136p33 is stabilized when *UL135* is disrupted). We previously generated a *UL135_stop_ UL136* intermediate virus that serves as the base BAC for the two new recombinant viruses created in this study (86). The “*UL136_myc_*” and “*UL136_myc_*Δ*K*→*R*” sequence mutations were polymerase chain reaction (PCR) amplified off of BACs previously made and characterized with these desired mutations using the following primers: UL138 (139bp) Rev and UL135 (854 bp) Fwd (43, 45). The sequences for the primers used for this amplification are listed in Table 1 below. The PCR amplified products were then recombined into the *UL135_stop_ UL136* BAC as described previously (43, 86). BAC integrity was tested by enzyme digest fragment analysis and sequencing of the *UL136* region using the same primers described in Table 1. All BAC genomes were maintained in SW102 *E. coli* and viral stocks were propagated by transfecting 15-20 μg of each BAC genome, along with 2 μg of a plasmid encoding *UL82* (pp71) into 5 x 10^6^ MRC-5 fibroblasts, allowed to propagate in MRC-5 fibroblasts and then subsequently purified and stored as previously described to generate the parent (P0) virus (39). To grow the next generation of each virus (P1), the infectious inoculum from the P0 virus was added to MRC-5 fibroblasts that were passaged into roller bottles and cultured for anywhere from 8-30 days depending on the recombinant virus being made. Viruses were subsequently purified and stored as previously described (39). Virus titers were determined by 50% tissue culture infectious dose (TCID_50_) on MRC-5 fibroblasts.

**Table 1.**
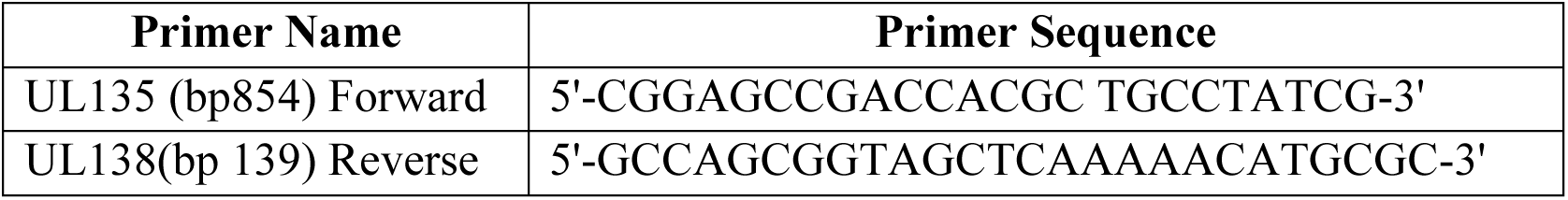
Primers used in this study for BAC recombineering.

**Table 2.** List of Antibodies used in this study.

| Antigen | Antibody | Concentration | Animal | Source |
| --- | --- | --- | --- | --- |
| myc epitope-M | 9B11 | 1:1,000 | Mouse | Cell Signaling |
| myc epitope-R | 71D10 | 1:1,000 | Rabbit | Cell Signaling |
| myc epitope-R |  | 1:1,000 | Rabbit | Proteintech |
| alpha-tubulin | DM1A | 1:2,000 | Mouse | Sigma |
| UL138 |  | 1:500 | Rabbit | Open Biosystems |
| UL135 |  | use at 2ug/ml | Rabbit | Open Biosystems |
| IE1&2 | 3H4 | 1:250 | Mouse | Gift, Dr. Thomas Shenk through John Purdy |
| pp150 |  | 1:15 | Mouse | Gift from Dr. William Britt |
| pp28 (UL99) | 10B4-29 | 1:50 | Mouse | Gift, Dr. Thomas Shenk |
| UL44 | 10D8 | 1:12,000 | Mouse | Virusys |

### Whole Genome Next-Generation Sequencing and Computational Analysis

Seven recombinant Human Cytomegalovirus Bacterial Artificial Chromosome (BAC) genomes DNA were extracted from SW102 *E. coli* propagating each HCMV genome using a BAC miniprep DNA isolation method. DNA was then sent to SeqCenter, LLC for short read Illumina whole genome sequencing using an Illumina NextSeq 2000 (200 Mbp sequencing package - https://.seqcenter.com/dna-sequencing/). Demultiplexing, quality control and adapter trimming was performed with bcl-convert (v3.9.3). The SNPs were identified using a python package called SNIPPY (https://github.com/tseemann/snippy, Accessed June 23 2022).

It’s based on BWA-mem/freebayes pipeline that identifies the mutations between a haploid reference genome and the next generation sequence (NGS) reads. The package was implemented with default parameters on University of Arizona High Performance Computing (HPC) environment, dedicating 80GB RAM for the job. SNIPPY was able to identify Single Nucleotide Polymorphisms (SNPs), Multi Nucleotide Polymorphisms (MNPs), Complex mutations as well as indels.

### Immunoblotting

Immunoblotting was performed as previously described (39, 43). Briefly, 50 μg of protein lysate were separated on 12% Bis-Tris gels by electrophoresis and transferred to 0.45um polyvinylidene di-fluoride (Immobilon-FL, Millipore) membranes. Proteins were detected using epitope or protein specific antibodies and fluorescently conjugated secondary antibodies using the Odyssey infrared imaging system (Li-Cor). All antibodies were used as listed in Table 2 below. Where indicated, cells were treated with 50-100μg of phosphonoacetic acid (PAA), 1-50μM MG132, or vehicle control. We are grateful to Dr. Thomas Shenk for the gift of antibodies from Drs. Thomas Shenk, William Britt, and John Purdy.

### Viral Growth Curves

Quantification of infectious virus produced by fibroblasts were determined by infecting MRC-5s at an MOI of 0.02 and subsequently collecting cells and medium over a 16-day infection time course. Virus titers were determined by TCID_50_ in MRC-5 fibroblasts.

### Quantification of Viral Genomes in Fibroblasts

Total DNA was isolated from ∼6x10^5^-7x10^5^ MRC-5 infected fibroblasts using Zymo Duet RNA/DNA isolation Kit (Zymo Research). Viral genomes were quantitated by quantitative PCR (qPCR) using the LightCycler 480 1X SYBR Green master mix (Roche) according to the manufacturer’s instructions. Primers specific to the non-coding b2.7 RNA gene were used. To determine the number of viral genomes present, viral DNA copy numbers were quantified relative to a BAC standard curve normalized to a cellular house-keeping gene RNase P as previously described (47).

### Infectious Centers Assay

CD34^+^ HPCs, isolated from human cord blood, were used to assess latency and reactivation of HCMV *in vitro* as previously described (34, 52). Briefly, CD34^+^ HPCs were infected at an MOI of 2 for 20 hours after which a pure population (>97%) of infected (GFP^+^) CD34^+^ cells were isolated via FACS (FACSAria, BD Biosciences Immunocytometry Systems, San Jose, CA) using a phycoerythrin-conjugated CD34 specific antibody (BD Biosciences) and propidium iodide to exclude dead cells. These cells were cultured in transwells above irradiated (4000 rads, ^137^Cs gammacell-40 irradiator type B, Atomic Energy of Canada LTD, Ottawa, Canada) M2-10B4 and Sl/Sl stromal cells for 10 days. The frequency of the production of infectious centers was measured using an extreme limiting dilution assay as described previously (34, 52). The frequency of infectious centers, based on the number of GFP^+^ cells at 14 days post plating, was calculated using ELDA, extreme limiting dilution analysis software (http://bioinf.wehi.edu.au/software/elda/) (87).

### Engraftment and Infection of huNSG Mice

All animal studies were carried out in strict accordance with the recommendations of the American Association for Accreditation of Laboratory Animal Care (AAALAC). The protocol was approved by the Institutional Animal Care and Use Committee (protocol 0922) at the Vaccine and Gene Therapy Institute (OHSU). NOD-*scid* IL2Rγ_c_^null^ mice were maintained at a pathogen free facility at Oregon Health and Science University in accordance with procedures approved by the Institutional Animal Care and Use Committee. Both sexes of animals were used. Humanized mice were generated as previously described (44, 88). The animals (12-14 weeks post-engraftment) were treated with 1 ml of 4% thioglycolate (Brewer’s medium; BD) via intraperitoneal (IP) injection to recruit monocytes/macrophages). After 24 h, mice were infected with HCMV TB40/E-*UL136myc* or *UL136myc* mutant-infected fibroblasts (approximately 10^5^ PFU per mouse) via IP injection. A control group of engrafted mice were mock infected using uninfected fibroblasts. The virus was reactivated as previously described (44, 88).

Data Availability. Raw viral genome sequences read have been deposited in the Sequence Read Archive (<u>SRA</u>) (BioProject ID: <u>PRJNA926319</u>).

### Quantitative PCR for Viral Genomes in HuNSG Mice

Total DNA was extracted from approximately 1mm^2^ sections of mouse spleen or liver using the DNAzol kit (Life Technologies), and processed as previously described (44). Primers and a probe recognizing HCMV UL141 were used to quantify HCMV genomes (probe =CGAGGGAGAGCAAGTT; forward primer = 5’ GATGTGGGCCGAGAATTATGA and reverse primer = 5’ATGGGCCAGGAGTGTGTCA). The reaction was initiated using TaqMan Fast Advanced Master Mix (Applied Biosystems) activated at 95°C for 10 minutes followed by 40 cycles (15s at 95°C and 1min at 60°C) using a StepOnePlus TaqMan PCR machine. Results were analyzed using ABI StepOne software.

## ACKNOWLEDGEMENTS

This study was supported by the National Institutes of Allergy and Infectious Diseases/ National Institutes of Health (NIAID/NIH) grants AI079059 and AI127335 to FG and AI127335 awarded to PC. We are grateful for the support of the Flow Cytometry and Human Immune Monitoring Shared Resource, supported through the Cancer Center Support Grant P30 CA023074 awarded to University of Arizona. We are grateful to Syed Shujaat Ali Zaidi (University of Arizona) for bioinformatic support and to Pierce Longmire for assistance with the SRA sequence submission. We thank Drs. James Alwine (University of Pennsylvania) and Lynn Enquist (Princeton University) for critical discussions on this work.

